# mTOR controls growth and internal architecture of human breast cancer spheroids

**DOI:** 10.1101/2024.02.24.580871

**Authors:** Katharina Hötte, Sabine C. Fischer, Alexander Schmitz, Michael Koch, Sanam Saeifar, Ernst H.K. Stelzer, Francesco Pampaloni

## Abstract

mTOR is a serine/threonine kinase participating in two distinct functional complexes: mTORC1 and mTORC2. mTORC1 regulates protein translation, cell cycle progression and autophagy. mTORC2 controls cell survival, proliferation and actin cytoskeleton reorganization. mTOR signaling is often deregulated during breast cancer. We investigated how mTOR and its downstream process autophagy determine the inner architecture of solid tumor aggregates at multiple levels. We focused on structures, which are morphologically altered during progression of cancer, including: nuclei number and volume, cell density, number, shape and volume of polarized structures. We developed a breast cancer spheroid model, in which the effects of mTOR and autophagy modulation were investigated by pharmacological inhibition mTOR signaling and induction autophagy, as well as using ATG7 KO spheroids. Spheroids were imaged with light sheet-based fluorescence microscopy and data were quantitatively analyzed. Our data suggest that mTORC1 regulates the spheroid volume, cell number and size. mTORC2 controls nuclei volume and determines growth and elongation of polarized acini inside a spheroid.

## Introduction

The mammalian target of rapamycin (mTOR) is an evolutionary conserved serine/threonine kinase. mTOR controls key signaling pathways such as cell survival, cell proliferation and autophagy, which are required for cellular and tissue homeostasis.

Two functionally distinct mTOR complexes are known. mTOR complex 1 (mTORC1) is responsive to cellular energy levels, i.e. ATP/AMP ratios, growth factors, and nutrient availability^1^. Growth factors act on mTORC1 through the PI3K–phosphoinositide-dependent kinase 1 (PDK1)–AKT pathway. The activation of mTORC1 via AKT triggers the direct phosphorylation of the p70 ribosomal protein S6 kinase (p70-S6K1)^2^ and inhibits the eIF4E-binding protein 1 (eIF4EBP1). Thereby, it promotes protein translation, and controls cell size and cell cycle progression^3^. Moreover, mTORC1 is involved in mRNA translation as well as in lipid synthesis and in the inhibition of autophagy under nutrient-rich conditions^4^. Fewer data are available on the function and regulation of mTOR complex 2 (mTORC2). Growth factors possibly activate mTORC2 via the PIK3 pathway, leading to full activation of AKT by phosphorylation of serine 473 (S473)^5^. However, mTORC2 is not responsive to nutrient availability^6^, and regulates cancer cell survival in an AKT-independent manner^7^. Through activation of the protein kinase C-alpha (PKCα), mTORC2 regulates cell migration and invasion by remodeling the actin cytoskeleton^8,9^, supposedly contributing to mammary morphogenesis in mice^10^.

Several morphological changes are associated with a typical cancerogenic phenotype, e.g. an increase in cell number and tissue volume, altered cell shape, reduced cell density^11^, increased nuclear size^12^, irregular nuclear shape^13^ as well as anomalous structures, e.g. inverted, apical-basal polarity^14^ or even a complete loss of polarization.

However, the contribution of mTOR to the development of such phenotypes is still unknown. Deregulation of mTOR is often found in cancer^15^. In breast cancer, the activation of mTOR correlates with a poor survival rate^16,17^. Therefore, mTOR is a promising target for therapies of breast cancer. Currently, mTOR inhibitors are tested in several clinical trials4. The most often studied mTOR inhibitors are rapamycin and its analogs, often referred to as rapalogs4. Rapamycin treatment inhibits mTORC1, leading to inhibition of p70-S6K1, as well as a rapid induction of autophagy.

We show that mTOR inhibition by rapamycin and nutrient starvation, as well as autophagy inhibition alters the growth and the morphology of a T47D breast cancer spheroid model. T47D cells show the typical molecular signature of luminal breast cancer, being oestrogen receptor positive (ER+), progesterone receptor positive (PR+) and human epidermal growth factor receptor 2 negative (HER2-)^18^. Multi-cellular tumor spheroids mimic several characteristics of tumors in vivo including cell-cell interactions, formation of polarized structures, and formation of a proliferative outer cell layer, a dormant layer and a necrotic core^19^. Since, mTOR signaling is differentially regulated in three-dimensional cell cultures^20^, we used small T47D spheroids composed of 2,000 seeded cells and an average diameter of 300 µm. These spheroids have no necrotic core, but develop a proliferating outer layer, a dormant region and form polarized structures. In the dormant region, we observed epithelial polarization during the formation of polarized acinar structures^21^. The usage of multi-cellular spheroids allowed us to study the effects of mTOR signaling in a more realistic tumor model. The cell-cell interactions and the formation of polarized structures, a process that naturally occurs in epithelial tissues, were analyzed quantitatively.

This study, which relies on image-based quantitative analysis of multi-cellular tumor spheroids, provides a better understanding of the pivotal role of mTOR signaling and autophagy during the establishment of a cancer phenotype in breast cancer. We found that mTOR and autophagy affect the tissue architecture of T47D cancer spheroids at multiple scales. We observed that the phenotype typical for cancerous tissues, e.g. reduced cell density^11^, increased nuclear size^12^ and loss of polarized structures^14^, depends on mTOR stimulation. Two groups of phenotypic changes, depending on the inhibitory stimulus of mTOR and autophagy, were identified. First, a reduction of the spheroid mass due to a decrease of cell size and cell number can be attributed to the inhibition of mTORC1. Second, a decrease in nuclear volume, which is accompanied by an increase in acinar volume and peri-acinar apoptosis. These phenotypic changes can be ascribed to the inhibition of mTORC2 by long-term rapamycin treatment. The acini show a less elongated, more spherical shape upon mTORC2 inhibition compared to the control, suggesting transformation to a less malignant phenotype.

Our data support mTOR as an effective target for luminal breast cancer, whose inhibition converts the spheroid morphology to a less aggressive phenotype. The morphological analysis approach described here could provide a groundbreaking tool for monitoring and predicting the outcome of rapalog treatments in breast cancer.

## Materials and Methods

### Cell culture and spheroid formation

T47D cells, purchased from American Type Culture Collection (ATCC), were cultured in RPMI1640 (Gibco) supplemented with 10% FBS (Gibco, Lot: 41Q6640K) and 2 mM L-glutamine (Gibco) at 37 °C, with 5% CO2.

Multi-cellular spheroids were generated with the liquid overlay technique^22^. Briefly, 50 µl of the cell suspension containing 2,000 cells were seeded in low melting agarose-coated (1% w/v) (Carl Roth) 96-well plates. The plates were incubated under standard cell culture conditions at 37°C and 5% CO2 in humidified incubators for 3 days to allow reproducible formation of spheroids. After spheroid formation, the cells were treated with Rapamycin (Sigma Aldrich), Bafilomycin A1 (Sigma Aldrich), 0.1% DMSO (Carl Roth) or Earle’s Balanced Salt Solution (EBSS) (Gibco) for further 7 days. The medium containing either Rapamycin, Bafilomycin, DMSO or EBSS was refreshed every three days.

### CRISPR Cas9 mediated ATG7 knock-out cell line

T47D ATG7 knock-out cells were generated with the Clustered Regularly Interspaced Short Palindromic Repeats (CRISPR)/Cas9 system by lentiviral transduction. The CRISPR/Cas9 system specifically targeting human ATG7 (K0142505) was purchased from Applied Biological Materials (ABM) Inc. (Richmond, Canada). For production of virus particles, 1x 106 HEK293T cells (purchased from ATCC) were transfected with 10 μg pATG7KO, 6.5 μg pCMVΔR8.91 and 3.5 μg pMD2.G. Plasmids were diluted in 450 μl water, and 50 μl 2.5 M CaCl2 solution were added. 2x HBS was added drop-wise to the DNA/CaCl2-solution, and incubated at room temperature for 20 minutes. The medium (DMEM supplemented with 10% FBS and 4 mM L-glutamine) was replaced by medium including 25 μM chloroquine. The DNA-mixture was added drop-wise to the cells, and incubated for 12 hours in the incubator. Virus particles were collected by harvesting the supernatant at 24, 48, and 72 hours after transfection. The virus particles were stored at -80°C for 4 days, before transduction. For transduction, T47D cells were seeded at 4x 105 cells per 6-well, and were transduced after 8 hours. 1 ml of viral supernatant and 1 ml of growth medium were loaded onto the cells. Polybrene was added to a final concentration of 8 μg/ml. Subsequently, the plate was centrifuged at 1.000x rcf for 1 hour at room temperature. The medium was refreshed after 12 hours. At day 3 following transduction, puromycin was added at a concentration of 2.5 μg/ml. After 4 days of puromycin treatment, cells were incubated in normal growth medium. T47D ATG7 knock-out cells were raised from single cell colonies.

### Quantitative analysis of the autophagy flux inT47D mRFP-GFP-LC3 spheroids

T47D cells, stably expressing the tandem fluorescence protein mRFP-GFP-LC3 (tfLC3) were used for monitoring the autophagy flux by wide field fluorescence microscopy. T47D-tfLC3 cells were produced as previously described by using the RCAS (replication-competent avian sarcoma leukosis virus long terminal repeat with splice acceptor) vector system^23,24^. Briefly, T47D cells were transiently transfected with the avian tumor virus receptor A (TVA) by using electroporation (preset “FF150” program, 4D-Nucleofector transfection system; Lonza, Cologne, Germany). The cell suspension was cultured with full medium at 37 °C and 5% CO2 for 24 hours. The medium was replaced by the viral stock supernatant, which contained the tfLC3 RCAS vector construct. The transfected cells expressed visible levels of mRFP-GFP-LC3 within ∼3 to 7 days. The transfected cells were sorted by fluorescence-activated cell sorting (FACS). For monitoring the autophagy flux of T47D-tfLC3 spheroids, spheroids were formed from 2,000 cells for 3 days in a 96-well plate, before treatment with indicated drugs. Spheroids were imaged for 72 hours with an 1 hour interval. Therefore, the Zeiss Cell Observer automated wide-field fluorescence microscope (Carl Zeiss AG, Oberkochen, Germany) equipped with incubator box and environmental control system (37 °C, 5% CO2), was used. The microscope was controlled by the Zeiss ZEN Blue software (version 1.1.1.0). For quantification of the autophagic flux with the mRFP-GFP-LC3 fluorescent sensor, it is assumed that the rate of change of the mRFP and GFP emission intensities is proportional to the rate of change of the number of autolysosomes and autophagosomes in the cells, respectively. Thus, the autophagic flux is expressed by plotting the ratio of the mRFP and GFP intensities over time.

### Immunofluorescence staining

Immunofluorescence staining of spheroids was performed as described^25^. Briefly, spheroids were fixed with 4% PFA for 20 minutes at RT. After washing with PBS, the spheroids were permeabilized with 0.3% Triton X-100 for 30 minutes. Spheroids were blocked for 3 hours at RT. Primary antibodies were diluted in blocking solution as follows: 1:200 beta-catenin (H-102, sc-7963), 1:100 GM130 (35, BD – 610822), 1:100 Caspase-3 (Asp175) (Cell Signaling - #9661). Spheroids were incubated with primary antibodies for 24 hours at 37°C and 300 rpm shaking in a Thermomix (Eppendorf). Spheroids were rinsed with PBS, before incubation with 1 µg/ml DAPI (Merck) and secondary antibodies (anti-mouse AF568: Molecular Probes - A10037, anti-rabbit Alexa Fluor 488 - Molecular Probes - A11008), 1:400 diluted in blocking solution for 24 hours at 37°C and 300 rpm shaking in a Thermomix. Spheroids were rinsed with PBS.

### Optical clearing and sample mounting

Spheroids were dehydrated with increasing concentrations of ethanol in deionized water (30%, 50%, 70%, 90%, 95%, 100%, 5 minutes each) at room temperature^25^. The dehydrated spheroids were transferred into BABB solution (1:2 v/v, benzylalcohol: benzyl benzoate) with a refractive index of 1.56, and incubated until onset of transparency. Samples were transferred into rectangular glass capillaries (Hilgenberg), which were previously cleaned with 1% Hellmanex (Hellma Analytics) in deionized water for 2 hours shaking on a Thermomix at 70°C. The capillaries were mounted onto a stainless-steel sample holders before being sequentially placed inside the microscope incubation chamber filled with thiodiethanol (TDE, Sigma) with a refractive index of 1.52.

### Image acquisition

Images were acquired with custom-built monolithic digital-scanned light sheet-based fluorescence microscopes (mDSLM, LSFM)^26^. The microscopes were equipped with an Epiplan-Neofluar 2.5x/0.06 illumination objective and a N-Achroplan 20x/0.5 detection objective (Carl Zeiss) and a Clara camera (ANDOR technology, Ireland). Three channels were acquired sequentially, with a z-spacing of 1.29 µm. Laser and filter sets: 561 nm laser, 607/70 bandpass filter; 488 nm laser, 525/50 bandpass filter; 405 nm laser, 447/55 bandpass filter.

### Image analysis

Raw image stacks of the nuclei channel were pre-processed in FIJI (ImageJ version 1.51d, Java version 1.6.0_24). The raw image stacks were cropped to the region of interest. The background intensity was subtracted from every slice using the function Subtract Background with a ball radius of 25 pixels. Noise was reduced by Gaussian filtering each slice with a radius of 1 pixel. Cell nuclei were extracted by means of an automated three-dimensional segmentation pipeline implemented in Wolfram Mathematica (version 10.2)^19^. The complete pipeline is available for download (http://www.physikalischebiologie.de/downloads). In brief, pre-processed image stacks were rescaled by a factor of 3.5 along z to obtain isotropic voxels. Hence, in the resized image, the voxels are isotropic with a pitch of 0.65 µm. Cell nuclei segmentation was achieved by local thresholding and a marker-controlled immersion-based watershed algorithm. The local threshold corresponded to the mean of the local intensity distribution in a range of ten voxels. Marker points for the watershed algorithm were detected by a multiscale Laplacian of Gaussian (LoG) blob detector with a minimal range of three and a maximum range of six voxels^27,28^. Marker points were increased by morphological dilation with a spherical structuring element and a radius of three voxels. Using the marker points as starting points, the final segmentation result was obtained by an immersion-based watershed algorithm^29^. Using Edelsbrunner’s algorithm ^30^, the cell nuclei centroids were used to compute an alpha shape with the alpha parameter set to 90 voxels. From the alpha shape, the approximate spheroid volume could be readily obtained. A cell graph^31,32^ was constructed with an edge distance threshold of 40 voxels capturing the spatial arrangement of cells and the local cell density within a spheroid. Features of all cell nuclei that had a volume greater than 150 voxels, and global features of the spheroid were stored in tabular format.

### Acini segmentation

Raw image stacks of the Golgi channel were pre-processed in FIJI (ImageJ version 1.51d, Java version 1.6.0_24). To obtain a homogeneous intensity distribution in the individual images of the z-stacks, the stacks were resliced. Each slice was preprocessed with the function Enhance Contrast with saturation 0.4 and setting normalize, and the resulting image stack was resliced again to obtain the original orientation. The subsequent acini detection was implemented in Wolfram Mathematica (version 10.4). The code is available for download (http://www.physikalischebiologie.de/downloads). The pre-processed image stacks were rescaled by a factor of 3.5 along z to obtain isotropic voxels and then scaled down by a factor of four. Hence, in the resized image, the voxels are isotropic with a pitch of 1.29 µm and the image was smoothed by a Gaussian filter with radius 5. To detect acinar regions in the spheroids, we applied minimum detection at multiple thresholds and combined the resulting images. For a given threshold h, extended minima were detected for which the range of intensities is not greater than h. We varied h between 0.03 and 0.14 and chose the most appropriate interval of h for each image manually.

The resulting binary image for the acinar structures was multiplied by a binary mask for the whole spheroid to discard outliers. We extracted the number of acini per spheroid and the volume of each acinus from the segmentation. Acini with a volume of less than 21.5 µm3 were discarded from the analysis. Finally, we calculated the elongation of an acinus as the ratio of the maximum distance of all voxels to the centroid and the radius of an equivalent ball with the same volume as the acinus.

### Caspase-3 cell segmentation

Raw image stacks of the caspase-3 channel were pre-processed in FIJI (ImageJ version 1.51d, Java version 1.6.0_24). The raw image stacks were cropped to the region of interest. The background intensity was subtracted using the function Subtract Background with a ball radius of 50 pixels. Caspase-3 positive cells were segmented with the function 3D Object Counter. Objects with a size of 150-10,000 voxels and an intensity threshold of 1,000 were segmented. We extracted the number of caspae-3-positive cells per spheroid and calculated the proportion of apoptotic cells to total cells per spheroid, based on the nuclei segmentation data.

### Western Blot

Spheroids from two 96-well plates per condition were lysed in a buffer containing 0.5% sodium deoxycholate, 1% NP-40, 0.1% sodium dodecyl sulfate, 1 mM EDTA in PBS, and freshly added protease inhibitors (Sigma-Aldrich) and phospho-stop (Roche). Spheroids were sonicated three times with an ultrasonic processor (UP50H, 50 W, 30 kHz) supplied by Hielscher ultrasound technology (Amplitude 0.6, and Cycle 1). Proteins were resolved on 12% acrylamide SDS-polyacrylamide gels, and transferred onto nitrocellulose membranes (GE Healthcare, Little Chalfont, UK). Membranes were blocked with fat-free dry milk in TBS-T. Primary antibodies against GAPDH (AM4300, Ambion), LC3 (NB100-2220, Novus Biologicals), P70-S6K1 and phophoP70-S6K1 (9202, 9206, Cell Signaling, Danvers), APG7 (Santa Cruz - sc-8668, n-20) were incubated over night at 4°C. Secondary horseradish peroxidase-conjugated antibodies (115-035-003, 111-035-003, Jackson Immuno Research, Newmarket, UK) were incubated for 1.5 hours at room temperature. Protein bands were visualized with an enhanced chemiluminescence detection reagent in the Chemocam documentation system (Intas, Goettingen, Germany).

### Statistics

We applied non-parametric statistics since the distributions of the measurements are not symmetric. For each measurement, we provide a Box-Whisker chart, in which the box represents the interquartile range, i.e. 50% of the data. The median is displayed as a horizontal line in the box. The whiskers indicate the minimum and the maximum of the data without outliers. Data points larger than 1.5 times the interquartile range are considered as outliers (black dots), data points larger than 3.5 times the interquartile range are considered as far outliers (gray dots). For visualization purposes, the figures display normalized data.

To compare the measurements for all treatments with those for spheroids in DMSO, we performed hypothesis testing. We applied the Mann-Whitney test with the Bonferroni correction for multiple testing. The hypothesis testing was performed on the raw data. For the measurements nuclei volume, cell density and acinar elongation, we discarded significances for differences of less than 5%.

## Results

### Long-term rapamycin treatment and nutrient starvation induce autophagy

To investigate the effect of mTOR signaling and its down-stream process autophagy on the inner architecture of luminal breast cancer spheroids, we used two different approaches, (1) the pharmacological modulation of mTOR and autophagy activity and (2) the genetic disruption of the autophagy flux by ATG7 knock-out. For pharmacological modulation, we used rapamycin treatment and amino acid starvation with EBSS to inhibit mTOR signaling and to activate autophagy, as well as bafilomycin treatment for the inhibition of autophagic flux at a late stage. Additionally, we generated a T47D ATG7 knock-out (ATG7 KO) cell line for the complete inhibition of autophagy. Spheroids were formed for three days, before the spheroids were treated for seven days with the different autophagy modulators. Spheroids formed from ATG7 KO cells were grown in normal growth medium. Seven days after treatment, spheroids were immunostained against GM130 to visualize the cis-Golgi apparatus, as well as against β-catenin to stain the adherens junctions. Cell nuclei were counterstained with DAPI. After optical clearing, *in toto* imaging of spheroids was performed with light sheet-based fluorescence microscopy (Figure 1A).

**Figure 1:**
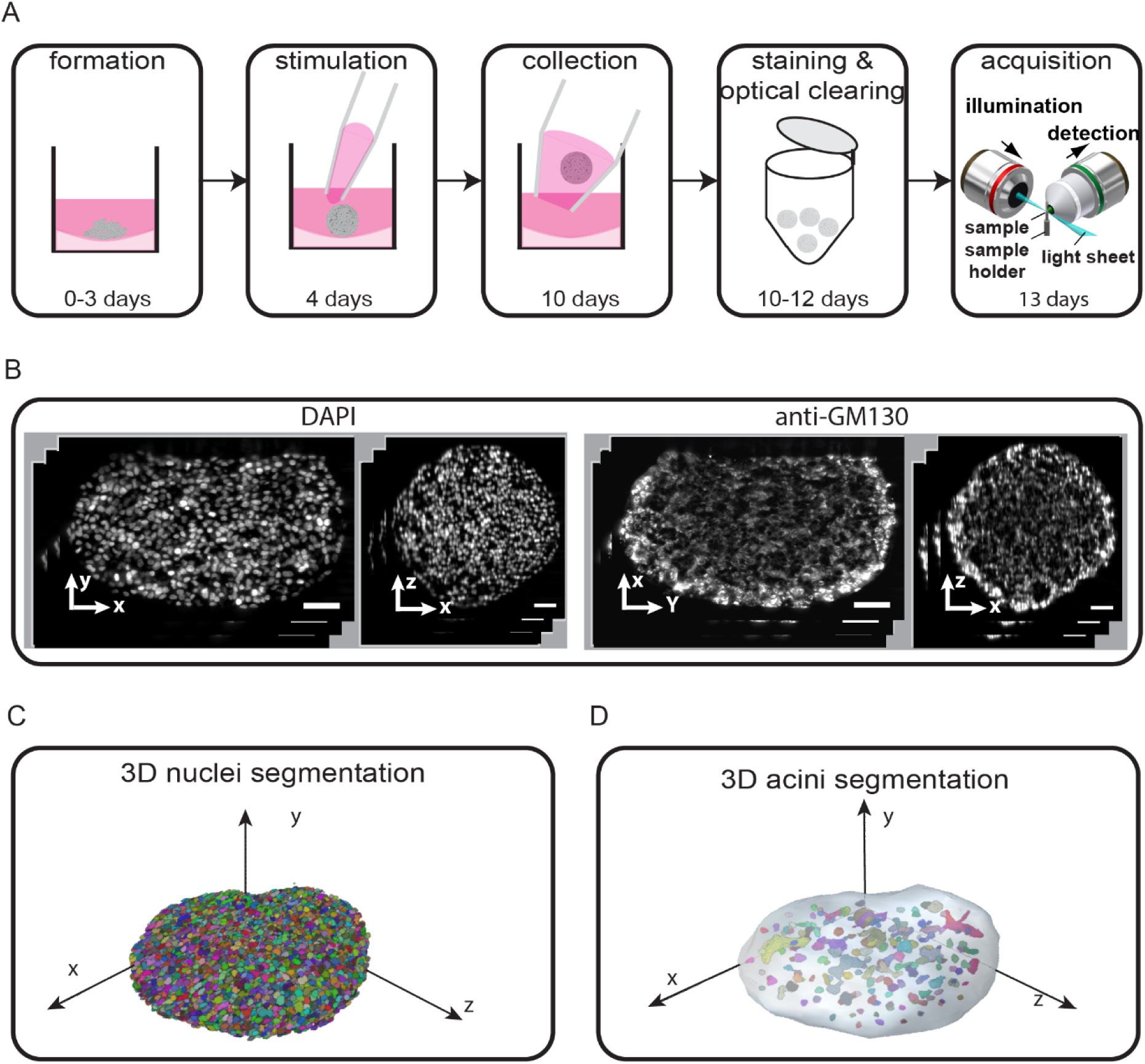
Schematic illustration of the experimental work flow. (A) Spheroids are formed for three days from 2,000 seeded T47D cells in 96-well plates. Each well is coated with 1% low-melting-point agarose. Starting on day four, spheroids are stimulated with autophagy-modulating drugs for seven days. Subsequently, they are collected for immunofluorescence staining. Prior to image acquisition with Light Sheet-based Fluorescence Microscopy (LSFM), spheroids are optically cleared with a Benzyl alcohol/Benzyl-Benzoate (BaBB) solution. (B) Single planes in x/y-and x/z-planes show cell nuclei (DAPI, left) and Golgi apparatuses (anti-GM130, right) in a representative T47D spheroid. Scale bar: 50 μm. Microscope: mDSLM. Excitation objective lens: 2.5x/NA0.06. Detection objective lens: 20x/NA0.5. GM130: 561 nm, bandpass filter 607/70. DAPI: 405 nm, bandpass filter 447/55. Scale bar: 50 μm. (C) Three-dimensional rendering of segmented cell nuclei. Each color represents a single segmented nucleus. (D) Three-dimensional rendering of segmented acini overlaid with spheroid surface (gray). Each color represents a single segmented acinus. The spheroid’s surface is approximated from the cell nuclei meshwork at the outer layer of the spheroid.

First, the efficiency of mTOR inhibition and, consequently, of autophagy induction by long-term treatment with rapamycin and nutrient starvation, as well as autophagic flux inhibition by bafilomycin treatment were quantified. Therefore, the autophagic flux was measured in T47D-tfLC3 spheroids over a time course of three days of treatment (Figure 2A and B). This assay allows the real-time observation of the autophagic flux, based on the fluorescence signal of monomeric green fluorescent protein (GFP) and red fluorescent protein (RFP), which are linked to LC3. GFP is more sensitive to the acidic pH of the lysosome compared to RFP and quenched during the formation of acidic autolysosomes^33^. For quantitative analyses of the autophagic flux, the ratios of the GFP/RFP signals were measured in the spheroids. A yellow signal represents the co-localization of GFP and RFP during autophagosome formation, while a red signal occurs during the formation of the autolysosomes. mTOR inhibition by rapamycin treatment and starvation leads to a strong red signal after 24 hours, which is even more notable after 72 hours (Figure 2A). These observations were confirmed by quantitative analyses of the GFP/RFP ratios. After six hours, the ratio decreases significantly, compared to the DMSO and bafilomycin treatments. This suggests that autophagy was strongly induced by starvation and 0.16 µM rapamycin treatment. The endpoint measurement after seven days of treatment was validated by Western blot analyses of the ratios between cytosolic LC3I and the lysosome membrane-bound LC3II (Figure 2C and D). Autophagy induction initiates an increase of LC3II level, which was determined by relative densitometry analysis. The average ratios of LC3II/LC3I upon rapamycin treatment was increased by a factor of five compared to the DMSO control (Figure 2C), while starvation increased the ratio by a factor of 2.8. Treatment with bafilomycin increased the LC3II to LC3I ratio by a factor of 2.2 (Figure 2D). These results indicate that both rapamycin treatment and starvation for seven days induced mTOR-mediated autophagy. The increase of the LC3II to LC3I ratio following bafilomycin treatment results from the accumulation of autophagosomes due to the inhibitory effect on the fusion with and acidification of lysosomes^34^. T47D ATG7 knock-out (KO) cells were generated with CRISPR/Cas9. Sequencing of the open reading frame and multiple alignment to the NCBI database showed a deletion of approximately 60 base pairs on allele 1 wild type and knock-out cells, suggesting that ATG7 on this allele is not functional in both cell lines (Figure S 1A). Multiple alignment the sequenced open reading frame of allele 2 revealed no mismatches in T47D wild type cells.

**Figure 2:**
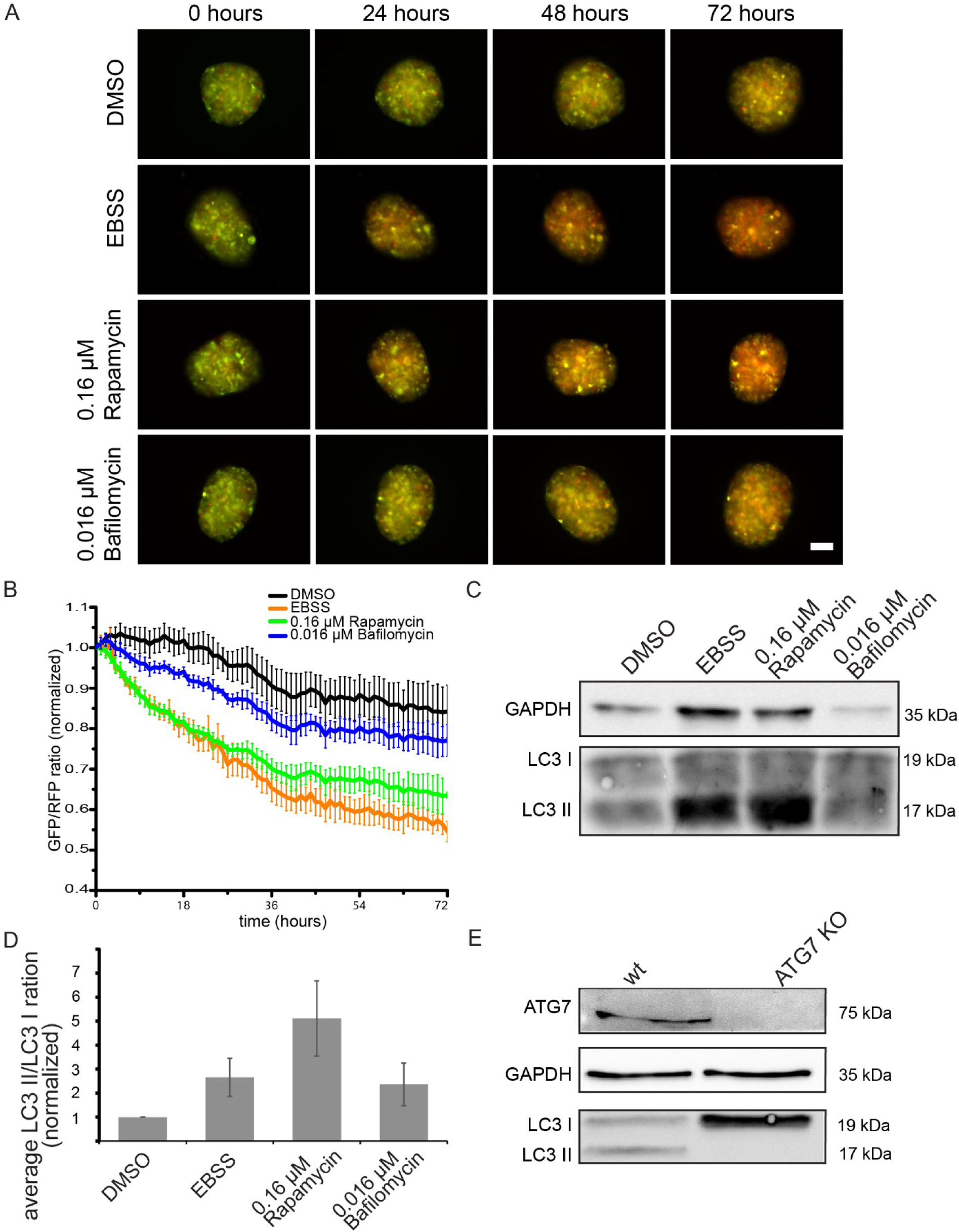
Monitoring the autophagy flux in T47D spheroids. (A) Images of T47D-tfLC3 spheroids 0, 24, 48 and 72 hours after treatment with rapamycin, bafilomycin and starvation (EBSS). (B) Quantitative evaluation of the GFP/RFP ration of T47D-tfLC3 spheroids over time. Data are normalized to GFP/RFP ratio at time point 0 hours. Error bars indicate SEM (n=18). (C) A representative image of a Western blot of T47D wild type spheroids after seven days of treatment. (D) Quantification of LC3 I to LC3 II turnover by relative densitometry. Data are normalized to the DMSO control. Error bars indicate SEM (n=4). (E) Western blot of T47D wild type and ATG7 knock-out spheroids. Microscope: Zeiss Cell Observer automated wide-field fluorescence microscope equipped with incubator box and environmental control system. (37 °C, 5% CO2). Images were recorded over 72 hours at an interval of 1 hour for each channel (GFP, mRFP) and each well.

However, two base pairs were deleted in ATG7 KO cells, leading two a frameshift and consequently the insertion of a stop codon (Figure S 2B). ATG7 KO spheroids were validated by Western blot analysis. Consistent with the absence of ATG7 in the knock-out spheroids, was the lack of the LC3II expression (Figure 2E). In contrast, the LC3I signal was clearly increased in the ATG7 KO spheroids, compared to wild type spheroids. These data indicate that T47D-ATG7 KO spheroids are incapable of performing autophagy. Thus, T47D-ATG7 KO spheroids are a suitable model to study the effects of autophagy inhibition on the inner architecture of breast cancer spheroids.

### mTOR modulation significantly changes the spheroid’s morphology

To explore the contribution of mTOR signaling to the formation of inner architecture in breast cancer spheroids, we first visually investigated global morphological changes in treated and ATG7 KO spheroids. As shown in figure 3, the volume of the spheroids treated with both rapamycin and EBSS starvation medium was reduced compared to the control group. In contrast, the volume of bafilomycin-treated and autophagy-deficient ATG7 KO spheroids was not affected (Figure 3). We found internal hollow lumina in spheroids at all conditions. The lumina are surrounded by well-aligned nuclei, suggesting the formation of polarized structures (acini) inside the spheroids, consistent with the luminal breast cancer origin of the T47D cells^21^. To confirm the polarization status of the acini, we analyzed the orientation of the cis-Golgi apparatus and of the cell nuclei with respect to the apical site of acinar structures (Figure 3, close ups). Independently from mTOR inhibition, inhibition of autophagy and knock-out of ATG7, most acini showed a reversed Golgi-nucleus orientation, similar to the phenotype found in the controls. In summary, these results suggest that mTOR inhibition influences the volume of the T47D breast cancer spheroids, while inhibition or deficiency of autophagy does not interfere with the spheroid size. The acinar polarity is unaffected by the inhibition of mTOR signaling and autophagy.

**Figure 3:**
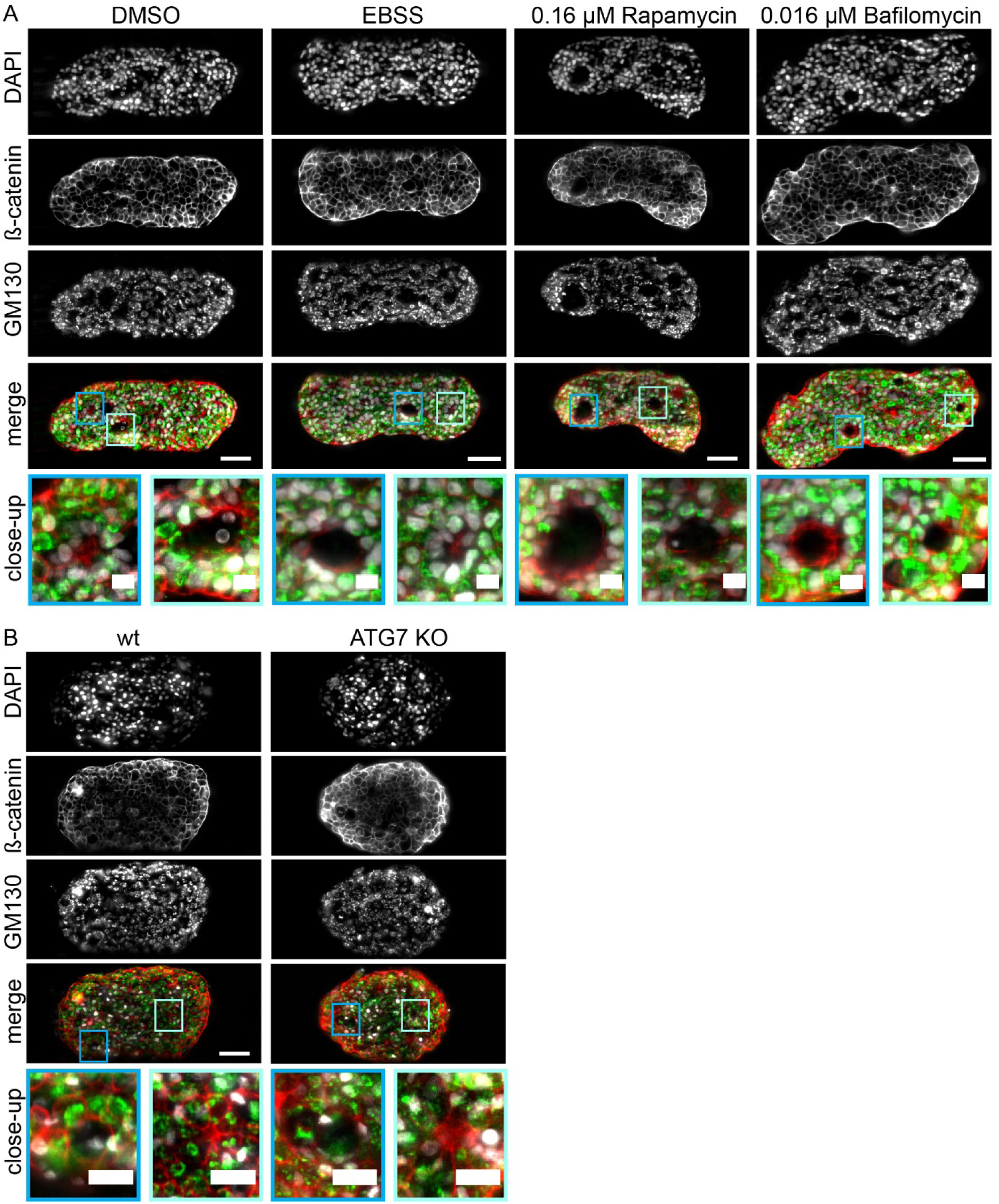
Long term mTOR inhibition alters morphology of breast cancer spheroids and acinar structures. (A) Single planes at the centers of optically cleared T47D spheroids after seven days of treatment with 0.1% DMSO as a control, EBSS starvation medium, 0.16 μM rapamycin and 0.016 µM bafilomycin. (B) Single planes at the centers of optically cleared T47D wild type and ATG7 knock-out spheroids. Cell nuclei (grey) were stained with DAPI, cell membranes (red) and Golgi apparatus (green) were visualized by immunofluorescence staining against β-catenin and GM130, respectively. Scale bar: 50 μm. Close-up (blue boxes) of acinar structures in the spheroids, either treated with rapamycin or DMSO. Scale bar: 20 μm. The prolonged induction of autophagy results in significantly decreased spheroid volumes compared to control spheroids. After rapamycin treatment, acinar structures within the spheroids are larger compared to those in control spheroids. Microscope: mDSLM. Excitation objective lens: 2.5x/NA0.06. Detection objective lens: 20x/NA 0.5. GM130: 561 nm, bandpass filter 607/70. β-catenin: 488 nm, bandpass filter 525/50. DAPI: 405 nm, bandpass filter 447/55.

### mTOR inhibition decreases spheroid volume and cell number in breast cancer spheroids

To dissect more precisely how mTOR signaling and its downstream process autophagy affect the architecture of breast cancer spheroids, we performed quantitative analysis of basic morphological parameters. To this aim, we applied a custom computational analyses pipeline^19^ for segmentation of cell nuclei in the entire three-dimensional spheroid. Using the acquired images of the DAPI-stained cell nuclei (Figure 4A, B and Figure S1), we performed segmentation of at least 15 spheroids per condition. The initial number of seeded cells, as well as the culture time of the spheroids, were the same for all conditions. Using the segmented nuclei data, we measured the following basic morphological parameters for each spheroid: (1) spheroid volume, (2) cell density, (3) number of cells and (4) nuclear volume.

**Figure 4:**
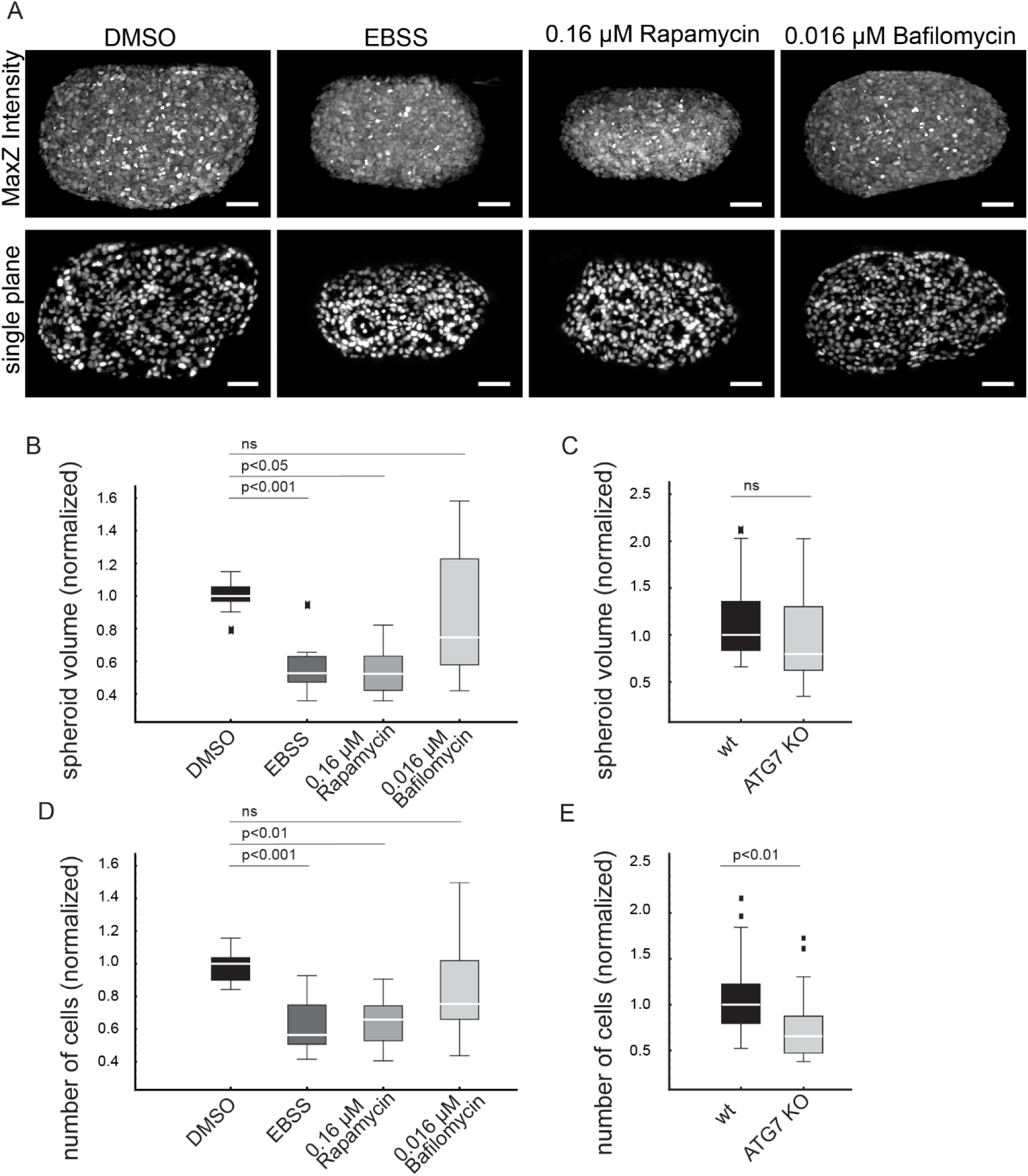
mTOR inhibition, but not autophagy inhibition reduces the spheroid volume. (A) Representative images of spheroids treated with 0.1% DMSO, EBSS starvation medium, rapamycin and bafilomycin. Maximum intensity projections show differences in spheroid volumes. MaxZ-intensity projections (upper panel) and single planes (lower panels) of a central slice show alterations of cell density in spheroids after mTOR inhibition. Boxplots of (B) normalized volumes of spheroids after long-term pharmacological mTOR and autophagy inhibition, (C) normalized volumes of wild type and ATG7 KO spheroids, (D) cell numbers of spheroids after long-term pharmacological mTOR and autophagy inhibition, (E) cell numbers of wild type and ATG7 KO spheroids. All data are normalized per batch. Autophagy induction for seven days by starvation with EBSS, and treatment with rapamycin, significantly reduces the number of cells in the spheroids. A prolonged induction of autophagy significantly reduces the volume of T47D spheroids. Prolonged autophagy inhibition by bafilomycin treatment or knock-out of ATG7 have no significant effect on cell number, average nuclear volume, local cell density, or spheroid volume, compared to controls. Boxplot parameters: box contains 50% of the data points, middle line of box is the median. Whiskers and outliers represent upper and lower 25% of the data. Outliers are outside 1.5x interquartile ranges. Statistics: Mann-Whitney test followed by Bonferroni correction (n=15).

Upon mTOR inhibition, induced with EBSS starvation medium, the median volume of spheroids was significantly reduced by 47%. A reduction of the median spheroid volume by 48% and 54%, respectively, was observed after the treatment with 0.16 µM and 0.08 µM rapamycin (Figure 4A, B and Figure S2A). This data indicates that mTOR inhibition significantly reduces the spheroid volume. In contrast, blocking the basal autophagic flux with bafilomycin (0.016 µM, 0.008 µM) did not affect the spheroid volume (Figure 4A, B and Figure S2A). The volume of ATG7 KO spheroids was slightly decreased (18%), compared to wild type spheroids, suggesting that autophagy inhibition has no significant impact on the regulation of spheroid volume.

### Autophagy controls the cell density of breast cancer spheroids

To identify the underlying cause of the volume reduction after mTOR inhibition, we quantified the number of cells per spheroid. The median number of cells after mTOR inhibition with 0.16 µM and 0.08 µM rapamycin, as well as through starvation with EBSS, was significantly reduced by 34%, 36% and 43%, respectively, compared to the control spheroids in 0.1% DMSO. The effect of mTOR inhibition by rapamycin treatment and starvation is well known to decrease the proliferation of T47D cells^35^, ^36^ via inhibition of the p70-S6K1 and 4E-BP1/eIF4E pathway^37^, and thereby leads to a reduction of spheroid volume. Autophagic flux inhibition with 0.016 µM or 0.008 µM bafilomycin did not show a significant influence on the cell numbers of the spheroids (Figure 4B, Figure S2B). Surprisingly, the median number of cells in ATG7 KO spheroids was significantly reduced by 35% (Figure 4E), compared to wild type spheroids, although no significant changes of the median spheroid volume were observed (Figure 4C). Based on these findings, we assumed that the spheroid volume is to a certain extent regulated by other processes besides the cell number. To assess this assumption, we compared the percentage of volume reduction and decrease in cell number. Interestingly, we found a difference of 14% and 18% after rapamycin (0.16 µM, 0.08 µM) treatment and 4% in starved spheroids (Table 1). In ATG7 KO spheroids, the difference between the percentage of spheroid volume and number of cells is 17%, suggesting changes in cell density after autophagy modulation. To pursue this rationale, we measured the local cell density in the spheroids, which is defined by the number of cells per unit of volume, and is dependent on the volume distribution of single cells within the spheroids. Both rapamycin treatment (0.16 µM and 0.08 µM) and starvation significantly increased the cell densities by 15%, 19% and 13%, respectively (Figure 5A, Figure S2C). In ATG7 KO spheroids, the cell density was significantly decreased by 18%, compared to wild type spheroids (Figure 5B). The cell density in bafilomycin-treated spheroids was not significantly altered (Figure 5A, Figure S2C). These data suggest, that mTOR signaling and, in particular, autophagy are involved in the regulation of the cell density.

**Figure 5:**
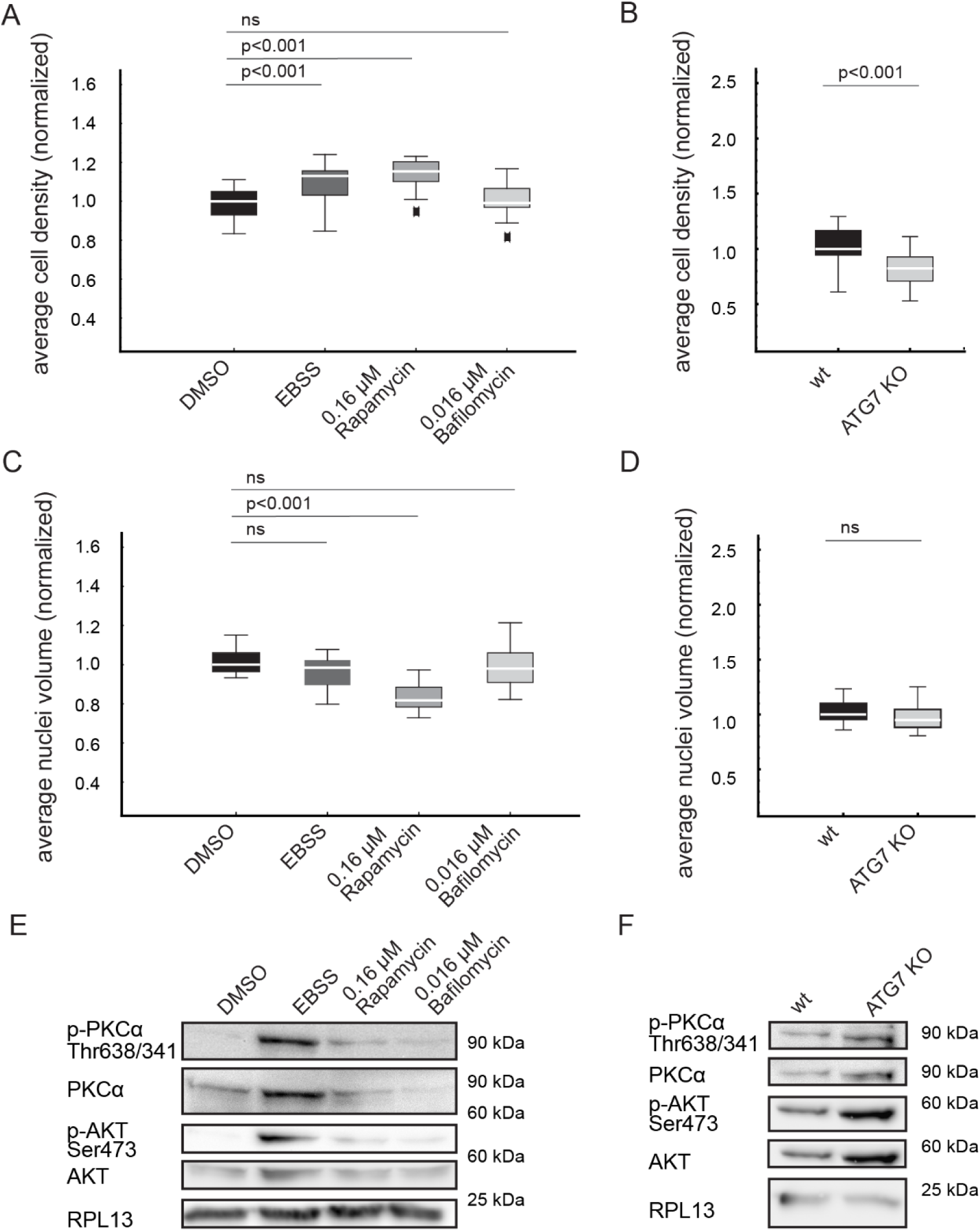
mTOR signaling controls the cell density and nuclear morphology in breast cancer spheroids. Boxplots of (A) cell density in spheroids after pharmacological modulation of mTOR signaling and autophagy, (B) cell density in ATG7 KO spheroids, (C) average nuclei volume of cells in spheroids after pharmacological modulation of mTOR signaling and autophagy, (D) average nuclei volume of cells in ATG7 KO spheroids. All data are normalized per batch. Long-term mTOR inhibition by starvation with EBSS and treatment with rapamycin significantly increases the density of cells in the spheroids. ATG7 knock-out decreases the cell density in spheroids, while bafilomycin treatment has no effect. Nuclear volume of rapamycin-treated spheroids is significantly decreased compared to DMSO controls. Boxplot parameters: box contains 50% of the data points, middle line of box is the median. Whiskers and outliers represent upper and lower 25% of the data. Outliers are outside 1.5x interquartile ranges. Statistics: Mann-Whitney test followed by Bonferroni correction (n=15). (E) Western blot analysis of Akt phosphorylation at S473 and PKCα phosphorylation at Thr638/341 in spheroids after pharmacological modulation of mTOR signalling and autophagy inhibition, as well as after ATG7 KO. RPL13 serves

**Table 1:** Spheroid volume is controlled by the number and density of cells.

| condition | median<br>spheroid<br>volume (%)<br>(normalized) | median<br>number of cells<br>(%)<br>(normalized) | difference<br>(spheroid<br>volume –<br>cell number) | cell density<br>(%)<br>(normalized) |
| --- | --- | --- | --- | --- |
| control | 100 | 100 | 0 | 100 |
| 0.16 μM<br>rapamycin | 52 | 66 | 18 | 115 |
| EBSS | 53 | 47 | 12 | 113 |
| 0.016 μM<br>bafilomycin | 75 | 75 | 50 | 99 |
| wild type | 100 | 100 | 0 | 100 |
| ATG7 KO | 82 | 65 | -11 | 82 |

Based on these data, we conclude that mTOR controls the spheroid volume at two different morphological scales: (1) the number of cells by regulating proliferation via p70-S6K1 and (2) the local cell density by regulating the cellular volume.

### mTOR inhibition by rapamycin, but not by starvation reduces the nuclei volume

We next asked whether mTOR signaling and autophagy are involved in the regulation of the nuclear morphology. For this approach, we measured the nuclei volume in rapamycin-, bafilomycin-treated, EBSS-starved, and ATG7 KO spheroids. Autophagy inhibition and ATG7 knock-out did not affect the nuclear volume. Surprisingly, rapamycin (0.16 µM and 0.08 µM) treatment, but not starvation, significantly reduced the nuclear volume by 19% and 16%, respectively, compared to the cell nuclei in control spheroids (Figure 5C, D, and Figure S2D). The data suggest that long-term mTOR inhibition by starvation and rapamycin treatment target distinct cellular processes. Previous studies showed that long-term rapamycin treatment (>24 hours), not only inhibits mTORC1, but also mTORC237. Therefore, we investigated the phosphorylation level of Akt at S473 and PKCα at Thr638/341, which are well-known to be phosphorylated upon mTORC2 activity as targets of mTORC2 signaling^38^. We observed an elevated expression and phosphorylation of Akt and PKCα after long-term starvation of T47D spheroids, compared to control spheroids. In spheroids, treated with rapamycin for seven days, the expression and phosphorylation level of PKCα and Akt was significantly reduced, compared to the phosphorylation levels after starvation (Figure 5E), indicating that mTORC2 is less active in rapamycin-treated compared to starved spheroids. These data confirm that starvation and rapamycin treatment target distinct cellular processes, probably via mTORC2 signaling. In ATG7 KO spheroids, the basal expression levels of Akt and PKCα are increased compared to wild type spheroids, while no effect is observed after autophagic flux inhibition with bafilomycin treatment.

### mTOR inhibition by rapamycin increases the acinar volume

To test whether mTOR signaling and its down-stream process autophagy are required for formation and development of polarized acinar structures in T47D breast cancer spheroids, we quantitatively determined (1) the number, (2) the volume, and (3) the shape of the acini inside the spheroids by three-dimensional segmentation based on the localization of the Golgi apparatus (Figure 1B, D).

First, we tested how mTOR and autophagy affect the number of acini in the spheroid (Figure 6A, B). For all conditions, we did not find any significant difference in the acini density. This indicates that mTOR and its downstream process autophagy do not affect the initial phase of acinar formation (Figure 6C, D, Figure S3B).

**Figure 6:**
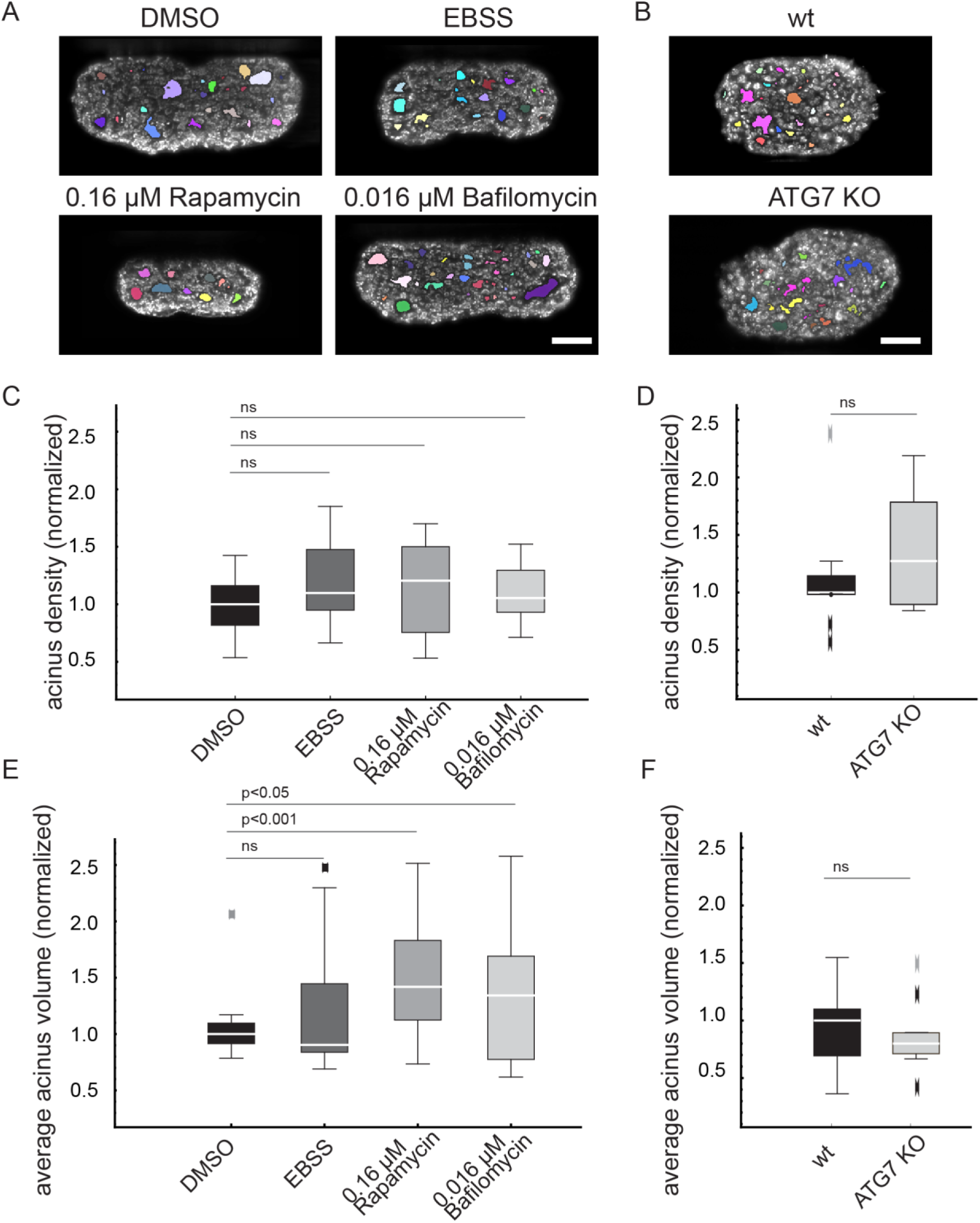
mTOR inhibition by prolonged rapamycin treatment increases acinar volume in T47D spheroids. Overlay of segmented acini (colored areas) and Golgi signal (grey) of single planes of optically cleared T47D spheroids (A) after pharmacological inhibition of mTOR and autophagy, as well as of (B) ATG7 KO spheroids. Scale bar: 50 μm. Boxplots of (C) the acinus density in spheroids after pharmacological mTOR and autophagy inhibition, (D) the acinus density in ATG7 KO spheroids (E) volume of acini in spheroids treated with rapamycin, bafilomycin, DMSO or starved with EBSS medium and (F) the volume of acinar structures in spheroids after ATG7 KO. Neither mTOR nor autophagy inhibition alter the density of acinar structures. The volume of acini in rapamycin treated spheroids is significantly increased, while starvation or inhibition of autophagy by bafilomycin or ATG7 KO do not interfere with the acinar volume. The Boxplot parameters: the box contains 50% of the data points, the middle line of the box is the median. Whiskers and outliers represent the upper and lower 25% of the data. Outliers are outside the 1.5x interquartile range. Statistics: Wilcoxon-Mann-Whitney-Test followed by Bonferroni correction (n=15).

Next, we investigated whether mTOR influences the subsequent acinar growth. To this aim, we measured the acinar volume. Interestingly, both mTOR inhibition by rapamycin (0.16 µM and 0.08 µM) and autophagy inhibition by bafilomycin (0.016 µM and 0.008 µM) led to a significant increase in acinar volume, by 42%, 57%, 34% and 32%, respectively. However, in ATG7 KO and starved spheroids the relative volume of acinar structures was not altered compared to control spheroids (Figure 6E, Figure S3). These data suggest that mTOR is involved in the development of acinar structures. Specifically, mTORC2 appears to control acinar growth as starvation-mediated mTOR inhibition, affecting mTORC1 only, did not alter the acinar volume. Additionally, these data show that the v-ATPase inhibitor bafilomycin, also affects the development of acinar structures.

We then analyzed the influence of mTOR-mediated autophagy on the elongation of acini towards pseudo-ductal structures (Figure 6D, Figure S3C). Following rapamycin treatment at 0.08 µM, acinar structures were significantly less elongated (6.1%), compared to the control group. At higher rapamycin concentrations (0.16 µM), acinar structures were less elongated (4.9%). This phenotype shows a more circular shape. No significant differences were observed for starved, ATG7 KO or bafilomycin-treated spheroids.

We conclude that inhibition of mTOR with rapamycin leads to an increase of the acinar volume. The data indicates that rapamycin-mediated, but not starvation-mediated mTOR inhibition controls the elongation of acinar structures. In summary, mTOR is not involved in early acinar development, but an autophagy-independent mTOR signaling pathway, supposedly encompassing mTORC2, is involved in acinar growth.

### mTOR inhibition increases apoptosis within the acini

Next, we investigated the interplay of mTOR and apoptosis, and the role of apoptosis in acinar formation. To test whether the increase in acinar volume is mediated by apoptosis we quantified the number of apoptotic cells, and monitored their spatial distribution after inhibition of mTOR and autophagy. The quantification of the amount of apoptotic cells was performed by staining cleaved caspase-3 positive cells in the spheroids. In general, apoptotic cells were rare in T47D spheroids, and they were mainly located in the center of small acinar structures (Figure 7A). In contrast, apoptotic cells found in larger acini, which predominantly formed after mTOR inhibition, were mainly localized in the acinar periphery.

**Figure 7:**
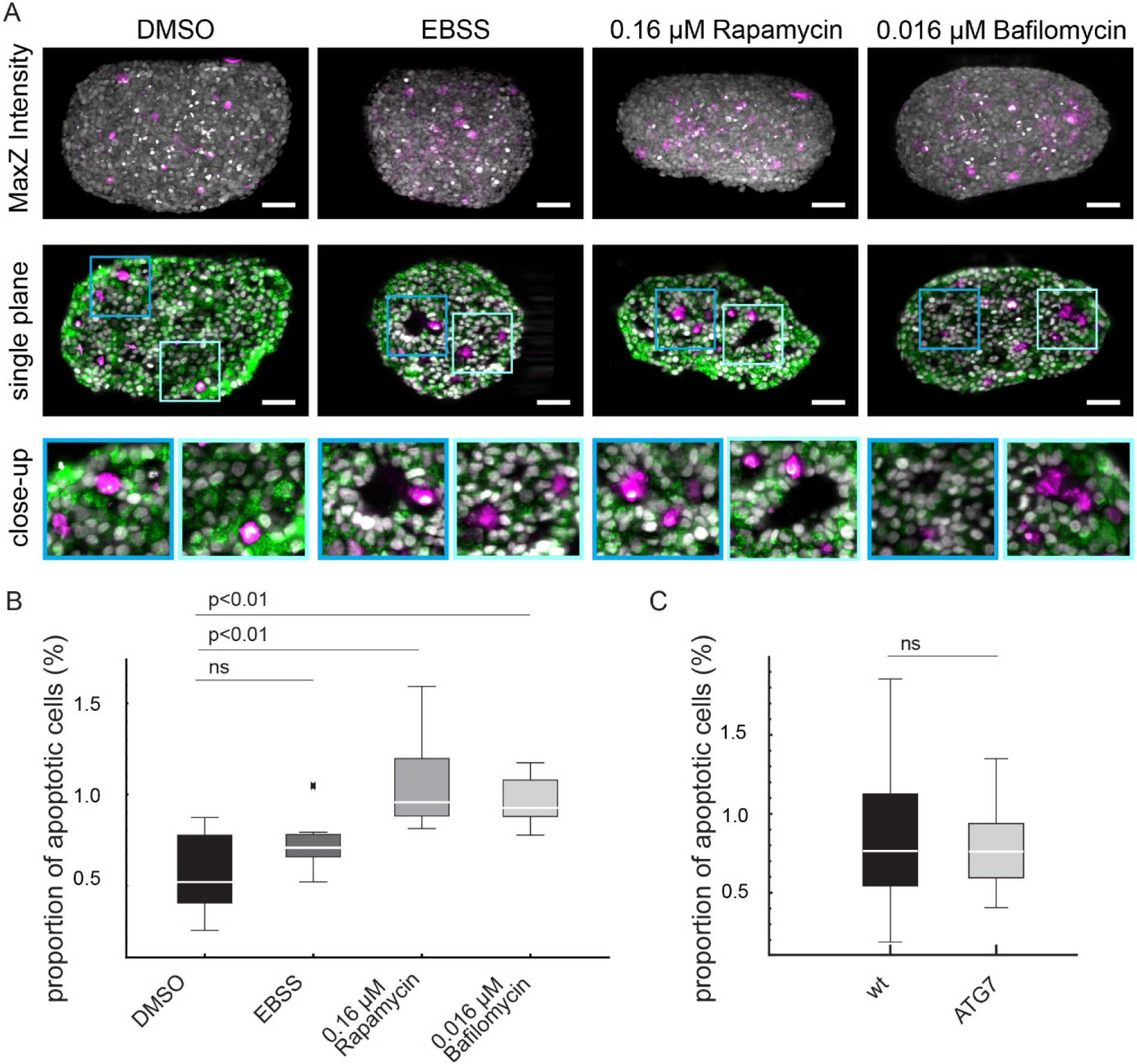
Long-term mTOR-mediated autophagy induced by rapamycin increases apoptosis in breast cancer spheroids. (A) Maximum intensity projections along the z-axis of optically cleared T47D spheroids after seven days of treatment with 0.1% DMSO, EBSS starvation medium, 0.16 µM rapamycin or 0.016 µM bafilomycin (upper panels), showing a summary of total nuclei (grey) and cleaved caspase-3 positive cells (magenta). Single planes show the spatial distribution of apoptotic cells (magenta), the Golgi apparatus (green), and nuclei (grey) in T47D spheroids (middle panels), seven days after treatment with 0.016 µM bafilomycin, 0.16 µM rapamycin, starvation with EBSS or 0.1% DMSO. Apoptotic cells and Golgi-apparatuses were detected by immunofluorescence staining against cleaved caspase-3 and GM130, respectively. Cell nuclei (grey) were counterstained with DAPI. Close-ups (lower panels) indicate that apoptotic cells occur predominantly in acini with smaller sizes. Scale bar: 50 μm. (B) Boxplot summarizing the ratio of apoptotic cells to the total cell number (n=15) in (B) spheroids after pharmacological modulation of mTOR signalling and autophagy, as well as in (C) spheroids of ATG7 KO cells. Boxplot parameters: the box contains 50% of the data points, the middle line of the box is the median. Whiskers and outliers represent the upper and lower 25% of the data. Outliers are outside the 1.5x interquartile range. Statistics: Mann-Whitney test followed by Bonferroni correction (n=12).

We found a significant higher number of apoptotic cells in rapamycin- and bafilomycin-treated spheroids compared to the control (Figure 7B). In rapamycin- and bafilomycin-treated spheroids, the proportion of apoptotic cells was twice as high as compared to the control (1% vs. 0.5% of the total number of cells). Starvation of spheroids and ATG7 KO did not alter the amount of apoptotic cells compared to the control (Figure 7B, C).

These results indicate that apoptosis is increased by rapamycin and bafilomycin treatment, but not by starvation or ATG7 KO. These data and the data obtained from quantitative analysis of the acinar volume, suggest that mainly apoptosis is required for the genesis of polarized structures in T47D spheroid. In small acini, apoptotic cells localize in the center, while in larger acini, apoptotic cells are mainly found at the acinar rim. This suggests that apoptosis is induced by mTOR in a starvation-independent process. Additionally, apoptosis is spatially regulated in T47D breast cancer spheroids, and contributes to the development of acinar structures.

## Discussion

In this study, we investigated the contribution of mTOR signaling and its downstream process autophagy on the morphology of breast cancer spheroids. This is of great interest, since mTOR is known to play a crucial role in the development and progression of breast cancer^39,40^. Following an approach similar to the one used by pathologists in the diagnosis of breast cancer^41,42^ we quantitatively analyzed volume, number of cells, cell density and size, nuclear volume, and luminal volume of polarized structures in the T47D spheroids (Figure 8). We found that mTOR regulates the inner architecture of T47D spheroids at multiple scales via its two distinct functional complexes: mTORC1 and mTORC2.

**Figure 8:**
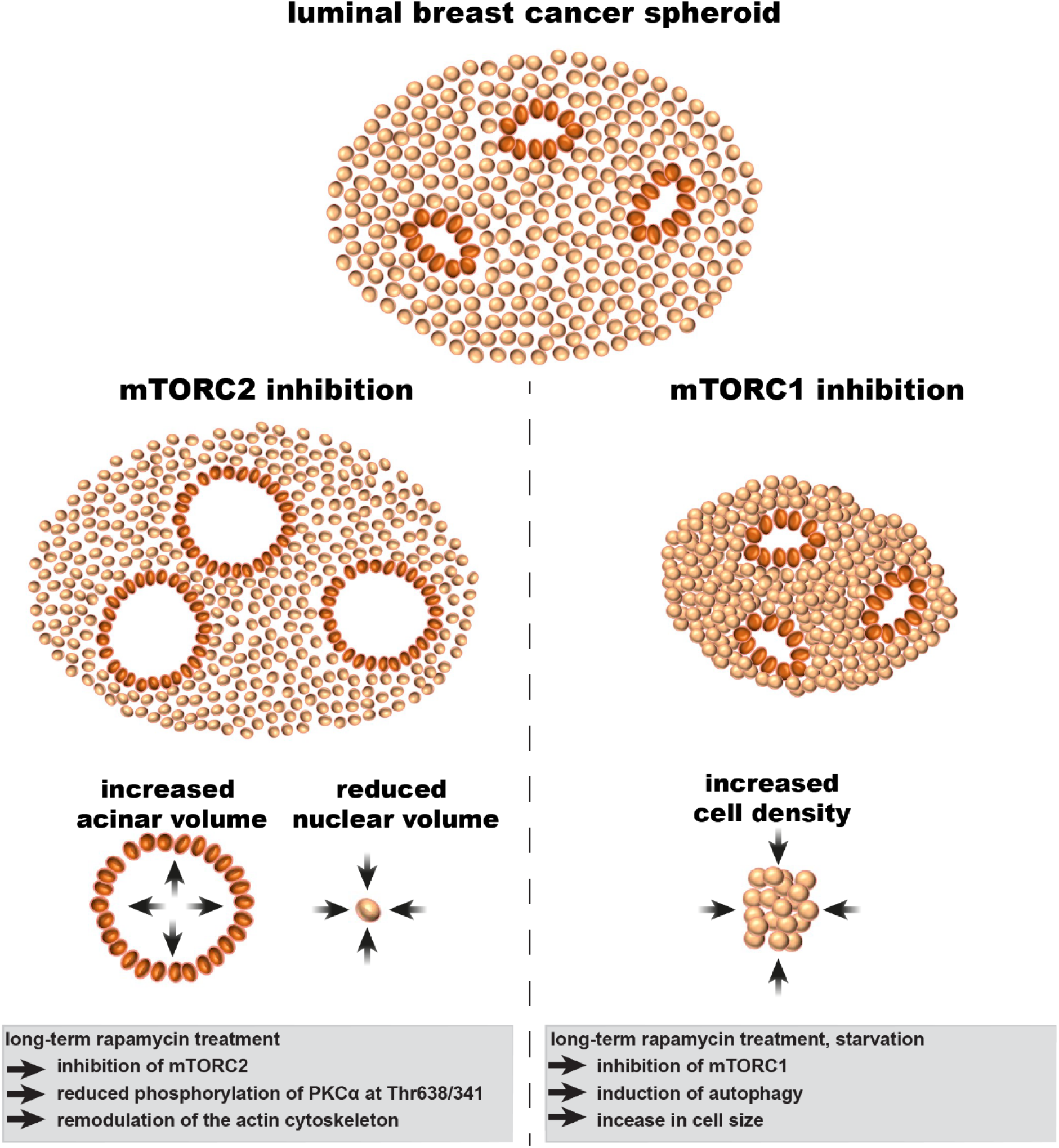
Different mTOR-mediated morphological changes of T47D spheroids as models for luminal breast cancer. Long-term rapamycin treatment for seven days inhibits mTOR in two distinct complexes, mTORC1 and mTORC2, while starvation only inhibits mTORC1. mTORC2 inhibition leads to a volume decrease of cell nuclei and a volume increase of acini, which are less elongated. This is probably mediated by reorganization of the actin cytoskeleton via PKCα signaling. Additionally, apoptosis is increased. Inhibition of mTORC1 leads to a reduction of spheroid volume and an increase in cell density, probably mediated by the inhibition of p70-S6K1, 4E-BP1 signaling and autophagy induction.

mTORC1 inhibition causes a significant shrinkage of breast cancer spheroid volume by reducing the number of cells. This was expected, since inhibition of mTOR, leads to downstream inactivation of p70-S6 kinase and eIF4E-binding protein 1, which are known to control cell cycle progression and proliferation^37^. Interestingly, in the autophagy-impaired spheroids formed with ATG7 KO cells, we observed no significant volume alterations, although the cell number was significantly reduced as well. We verified that autophagy, induced by starvation and rapamycin treatment, specifically regulates the spheroid volume by increasing the cell density, i.e. the spheroids become more compact under these conditions. In contrast, complete impairment of autophagy in ATG7 KO spheroids significantly decreased the cell density, i.e. the treated spheroids were less compact than the control. Our findings provide evidence that autophagy specifically regulates the tumor spheroid volume by reducing the cell size but not the cell number.

Alterations of cell size are often associated with changes of the nuclear volume. In pathological context, an increase in nuclear size correlates with increased proliferation and reduced survival rates of patients^12,43^. In tumor biology, alterations of nuclear shape and volume are reliable indicators for cancer progression^44^. However, the underlying mechanisms are poorly understood. We found that that mTOR inhibition by prolonged rapamycin treatment, but not by starvation, significantly decreased the nuclear volume in spheroids. As already pointed out, while nutrient starvation inhibits mTORC1 only, and triggers autophagy^45^, rapamycin inhibits both mTORC1 and mTORC2^46^, whereas mTORC2 inhibition onsets after prolonged (over days) rapamycin treatment^47^. Indeed, by measuring the basal expression and phosphorylation level of the downstream targets of mTORC2, namely Akt and PKCα, we showed that the activity of mTORC2 in long-term rapamycin-treated spheroids is strongly reduced compared to long-term starved spheroids. Therefore, we inferred that the decrease of nuclear volume does not depend on inhibition of mTORC1, but rather on inhibition of mTORC2. mTORC2 is known to induce re-organization of the actin cytoskeleton^48^. Since the actin cytoskeleton plays an important role in the regulation of nuclear morphology^49^, and loss of cortical actin has been shown to interfere with nuclear shape and integrity^50^, we hypothesize that a reduced nuclear volume after mTORC2 inhibition, however this hypothesis has yet to be substantiated by experimental data.

Loss of cell polarization and filling of the luminal space in mammary glands are critical structural alterations found in breast cancer^51^. In mammary cysts derived from a non-malignant mammary cell line (MCF10A), the development of acinar lumina is regulated by both, autophagy and apoptosis^52,53^. In the T47d spheroids, we observed that in breast cancer spheroids, neither inhibition of mTOR nor impairment of autophagy in ATG7 KO spheroids affected the acinar density (number of acini per spheroid). This suggests that mTOR signaling is not required for the initiation of acinar structures. In order to investigate the role of apoptosis and its interplay with mTOR during acinar development, we determined the number and spatial distribution of apoptotic cells in treated T47D spheroids. We observed that apoptosis is a rare event during maturation of T47D breast acini, consistently with what found in prostatic acini^54^. At variance with starvation and ATG7 KO, prolonged rapamycin treatment increased the number of apoptotic cells. Apoptotic cells were prevalently found proximal to the acini. In small acini, apoptotic cells localized in the luminal center, while in larger ones, apoptotic cells appeared in the polarized cell layer delimiting the lumen. Based on these data and previous studies, In summary, the initiation of acini formation is mediated by apoptosis only^55,56^. The extent of apoptosis can be modulated through mTORC2. In contrast, neither mTORC1 nor autophagy are involved in this process.

Elongation and growth of breast acini occur during physiological processes such as puberty and pregnancy^57^. In contrast, aberrations in the acinar volume, accompanied with apical-basal polarity are hallmarks of cancer^58,59^. To test whether the acinar phenotype can be modulated by mTOR, we quantified the volume and shape of acini in T47D spheroids. We measured an increase of acinar volume following bafilomycin and rapamycin treatment, while starvation and ATG7 KO did not affect the acinar volume, suggesting that autophagy plays only a minor role in acinar genesis. Despite we measured an increase of apoptosis after bafilomycin and rapamycin treatment, it is unlikely that apoptosis alone (∼1% of total cells), accounts for the acinar volume increase (∼50%), indicating that apoptosis is only marginally involved in this process. These data suggest that the increase of acinar volume is an mTORC2-mediated process. These data suggest that the increase of acinar volume is an mTORC2-mediated process. Prolonged rapamycin treatment resulted into a less elongated acinar shape, compared bafilomycin-treated, starved, or ATG7 KO spheroids These results are consistent with in vivo studies, showing that mTORC2 regulates the size and the branching of breast acini in mice^60^. Thus, both, the volume increase and the elongated shape of the acini, are mTORC2-mediated processes, possibly driven by rearrangements of the actin cytoskeleton via PKCα^9^.

In summary, we applied multiscale image analysis to study the role of mTOR and its downstream process autophagy on the morphogenesis of breast cancer spheroids. We show that mTOR regulates cellular phenotypes that are often altered during the progression of cancer. This includes the overall size of the spheroid, the number of cells, the development of polarized structures, the cell density and the nuclear size. Further, we collected evidence pointing to the distinct roles of mTORC1 and mTORC2 in the regulation of spheroid morphology. As summarized in (Figure 8), we found that mTORC1 is required for the regulation of cell growth and cell size, while mTORC2 is implicated in the regulation of nuclear morphology, as well as the elongation and growth of polarized structures. Moreover, the data indicate that mTOR is involved in the regulation of apoptosis in the peri-acinar region.

## Author contribution

K.H. and F.P. designed the study. K.H. M.K. and S.S. performed the experiments. A.S., S.C.F. and K.H. performed the data analysis. S.C.F. and KH performed statistical analysis. K.H., F.P. and E.H.K.S. wrote the manuscript. All the authors read and commented the manuscript.

## Supplements

**Figure S3:**
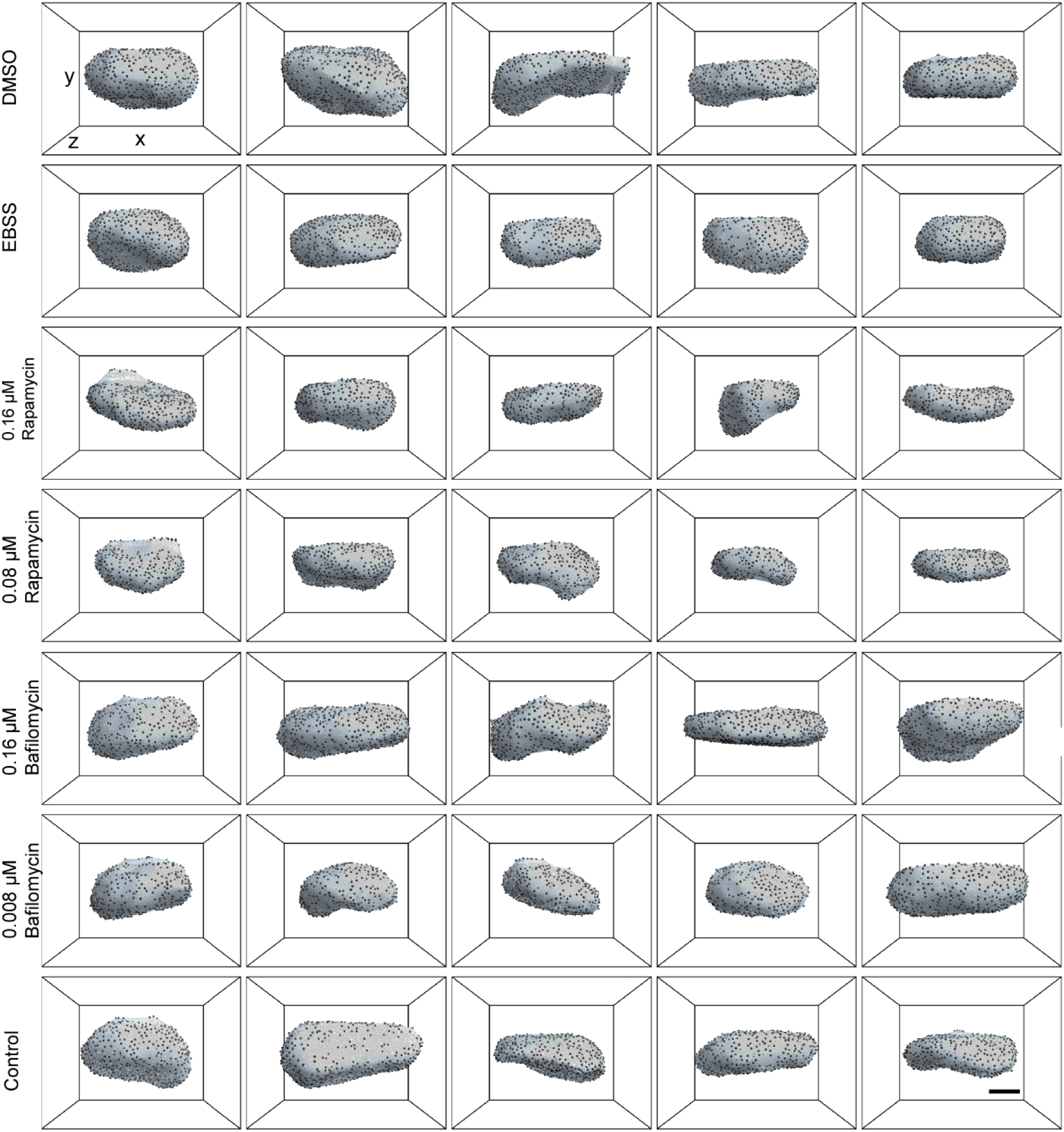
Overview of the surface approximations of T47D spheroids. Five representative spheroids for each condition are shown as three-dimensional surface approximations. The surfaces are calculated by interpolations of the meshwork formed by the outer nuclei layer. Scale bar: 50 µm.

**Figure S4:**
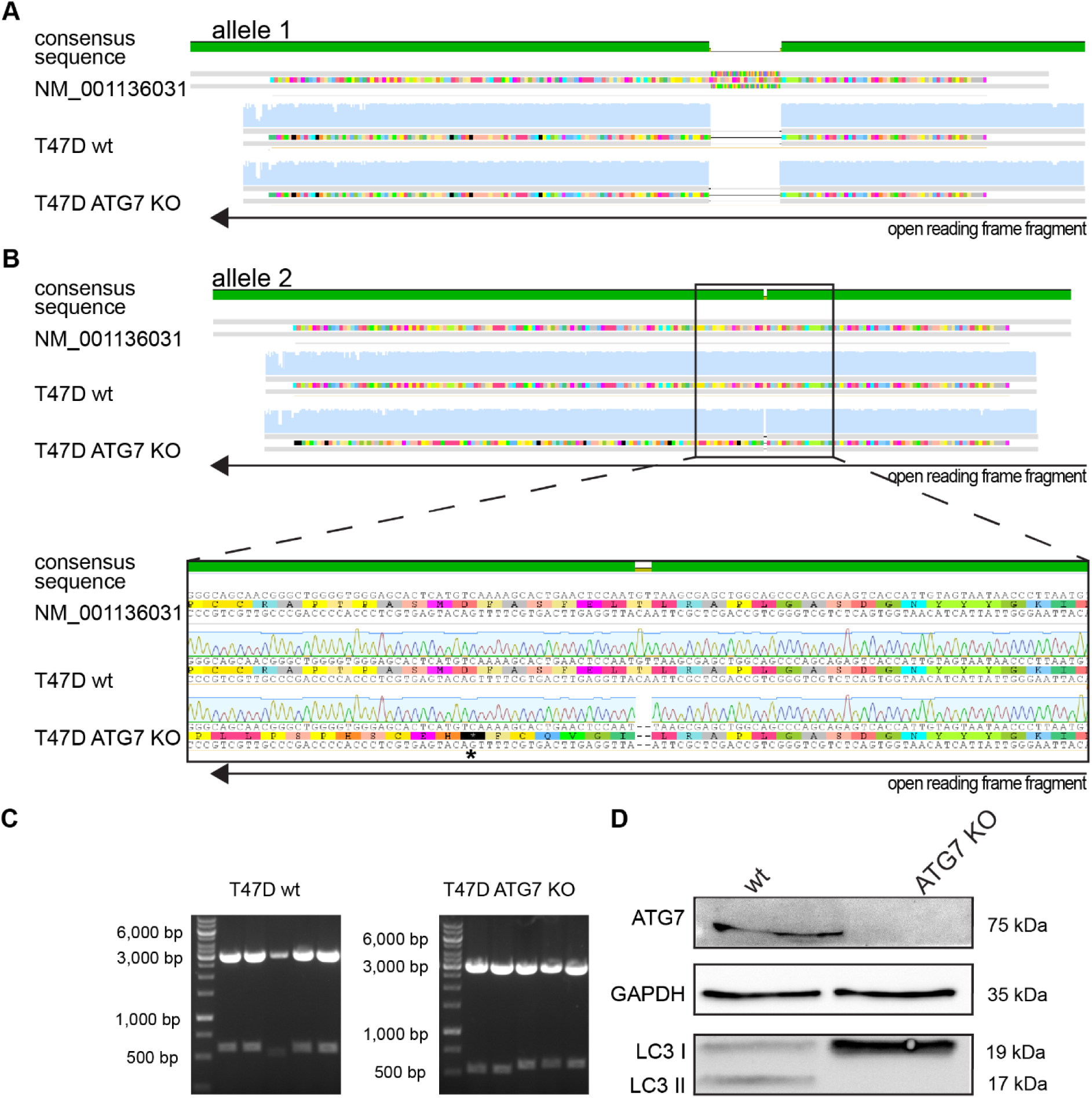
CRISPR/Cas9 mediated ATG7 knock-out leads to functional loss of autophagy. (A) Alignment of the Sanger sequencing results of the human atg7 gene on allele 1 of T47D wild type and ATG7 knock-out cells compared to the NCBI entry (NM_001136031). The green bar indicates the consensus sequence and the black arrow indicates the reading direction of the open reading frame fragment. The sequencing results reveal a deletion of approximately 60 base pairs on allele 1 in T47D wild type and ATG7 knock-out cells. (B) Alignment of Sanger sequencing results of the human agt7 gene on allele 2 of T47D wild type and ATG7 knock-out cells compared to the NCBI entry. The consensus sequence shows a mismatch of two base pairs, leading to a pre-mature stop codon (asterix) downstream of the open reading frame that is caused by a frame shift. (C) Agarose gel electrophoresis of the ATG7 PCR product (∼500 bp) cloned into the pGEMTEasy backbone (∼3,000 bp) after enzymatic restriction. (D) Western blot analysis of wild type (wt) and ATG7 knock-out (ATG7 KO) cells, verify the functional loss of autophagy, by detecting the ATG7 protein expression. No LC3 II is detected in ATG7 KO cells. GAPDH serves as a loading control.

**Figure S5:**
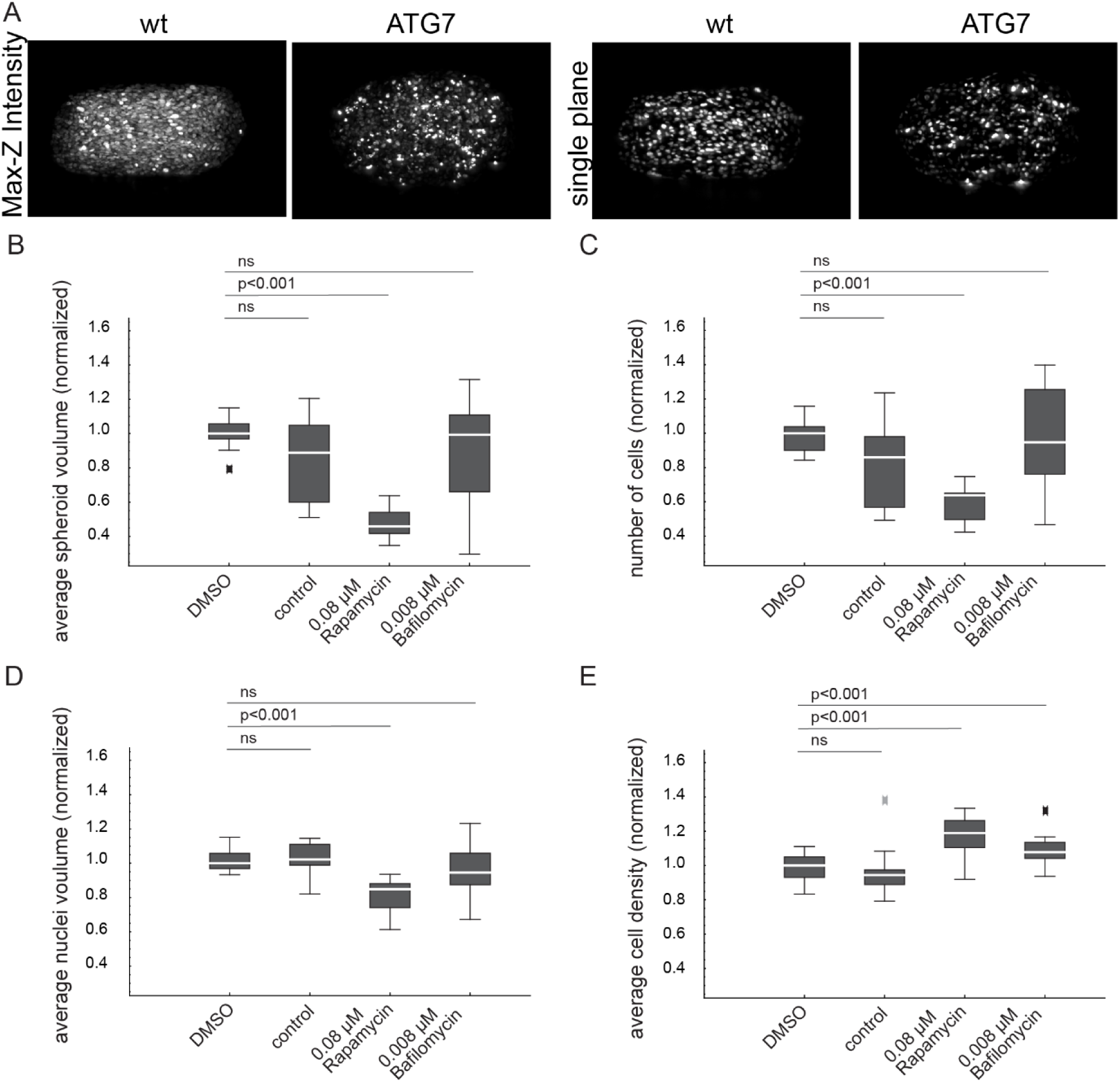
Long-term autophagy induction increases cell density and reduces nuclear volume in breast cancer spheroids. (A) Max-Z intensity projections and single planes of wt and ATG7 KO spheroids. Boxplots of (B) cell numbers, (C) average nucleus volume, (D) average local cell density, and (E) spheroid volume in T47D spheroids treated with 0.1% DMSO or normal growth medium as controls, as well as 0.08 µM rapamycin and 0.008µM bafilomycin. All data are normalized per batch. Autophagy induction for seven days, by treatment with 0.08 µM rapamycin significantly reduces the number of cells in the spheroids. The local cell density is significantly increased after treatment with 0.08µM rapamycin and 0.008 µM bafilomycin. The nuclear volume of rapamycin-treated spheroids is significantly decreased compared to DMSO and growth medium controls. Prolonged induction of autophagy significantly reduces the volume of T47D spheroids. Prolonged autophagy inhibition by bafilomycin treatment has no significant effect on cell numbers, average nuclear volume or spheroid volume, compared to both controls. Boxplot parameters: the box contains 50% of the data points, the middle line of the box is the median. Whiskers and outliers represent the upper and lower 25% of the data. Outliers are outside the 1.5x interquartile range. Statistics: Mann-Whitney test followed by Bonferroni correction (n=15).

**Figure S6:**
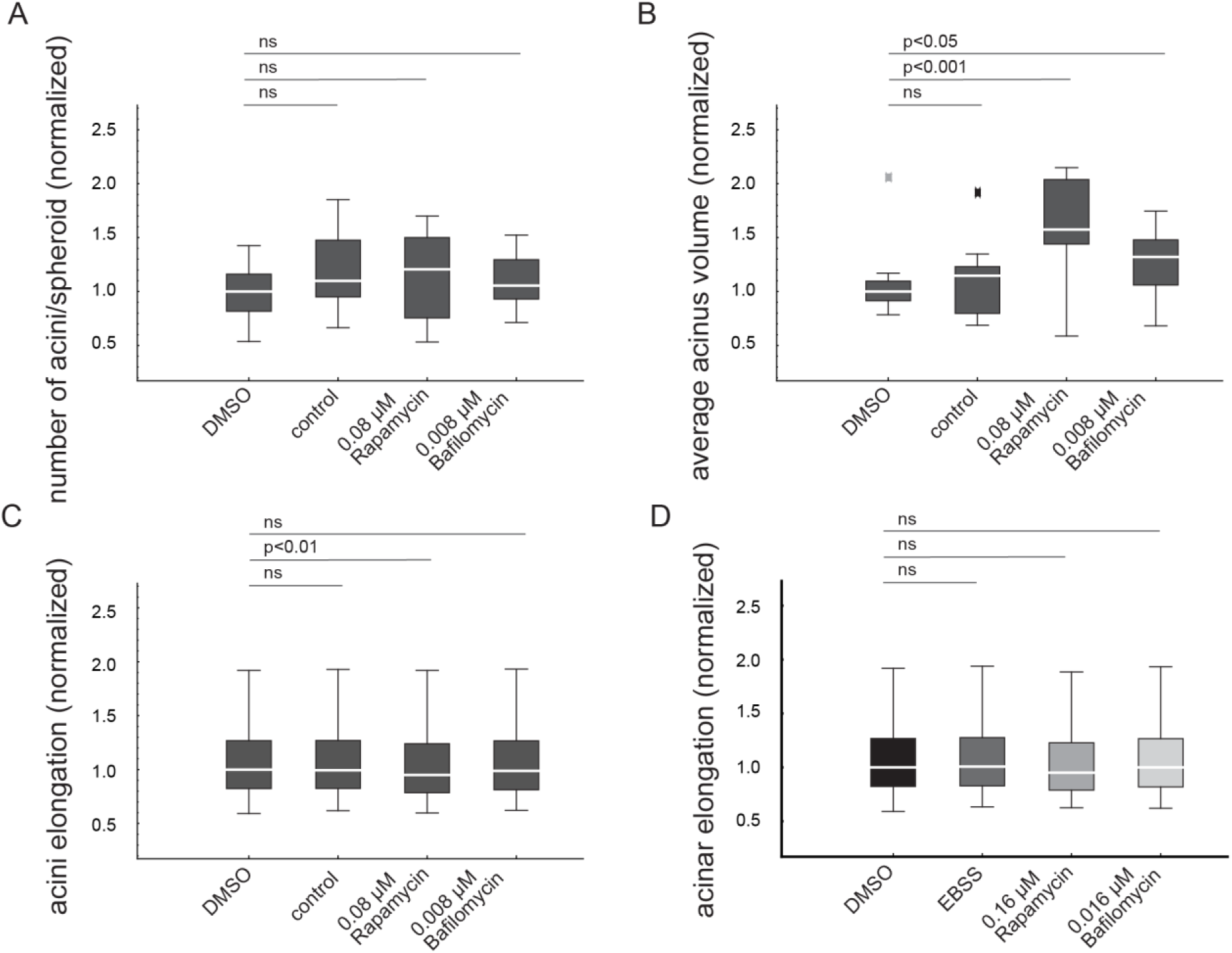
Autophagy inhibition and induction increase the acinar volume in breast cancer spheroids. T47D spheroids treated with 0.1% DMSO or normal growth medium as controls, as well as 0.08 µM rapamycin and 0.008µM bafilomycin. Acini were segmented in three dimensions based on the Golgi staining. The resulting segmentation is overlaid in red. Scale bar: 50 μm. Boxplots of (A) acini numbers, (B) acinar volume and (C, D) acini elongation in spheroids treated with rapamycin, bafilomycin, EBSS, DMSO or growth medium. All data are normalized per batch. Autophagy inhibition by 0.008 µM bafilomycin does not affect the number or elongation of acini in T47D spheroids compared to the controls. The acinar volume in these spheroids, however, is significantly increased. Boxplot parameters: the box contains 50% of the data points, the middle line of the box is the median. Whiskers and outliers represent the upper and lower 25% of the data. Outliers are outside the 1.5x interquartile range. Statistics: Mann-Whitney test followed by Bonferroni correction (n=15).

